# A transgene harboring the human Glucose Transporter1 (GLUT1) gene locus ameliorates disease in GLUT1 deficiency syndrome model mice

**DOI:** 10.1101/2025.04.05.647372

**Authors:** Maoxue Tang, Sasa Teng, Ashley Y. Kim, Yueqing Peng, Umrao R. Monani

## Abstract

Proper brain function relies on an adequate supply of energy – mainly glucose – to power neuronal activity. Delivery of this nutrient to the neuropil is mediated by the Glucose Transporter1 (GLUT1) protein. Perturbing glucose supply to the brain is profoundly damaging and exemplified by the neurodevelopmental disorder, GLUT1 deficiency syndrome (GLUT1DS). Resulting from haploinsufficiency of the *SLC2A1 (GLUT1)* gene, GLUT1DS is characterized by intractable infantile-onset seizures and a disabling movement disorder. Ketogenic diets, which supply the brain with an alternate energy source, ketone bodies, are currently the preferred therapeutic option for Glut1DS patients but do not address the underlying cause – low brain glucose – of the disease. One intuitively appealing therapeutic strategy that does, involves restoring GLUT1 levels to the patient brain. Here, we demonstrate that transgenic expression of the human *GLUT1* genomic locus in a mouse model of GLUT1DS raises brain GLUT1 levels and reduces disease burden. Augmenting GLUT1 levels in mutants correspondingly raised cerebrospinal fluid (CSF) glucose levels, improved motor performance and reduced the frequency of seizures characteristically observed in GLUT1DS. Interestingly, the increased GLUT1 in mutants harboring the human *GLUT1* locus was at least partly the result of an increase in murine *Slc2a1 (Glut1)* activity, most likely the effect of a long non-coding RNA (lncRNA) embedded in the human transgene. Collectively, our work has not only shown that repleting human GLUT1 mitigates GLUT1DS but also has yielded transgenic mice that constitute a useful tool to test and optimize clinically promising agents designed to stimulate this gene for therapeutic purposes.

## Introduction

The human brain consumes a disproportionate amount of an individual’s total energy intake. Indeed, while it constitutes just 2% of the weight of a typical adult human, the brain accounts for roughly a quarter of the body’s energy requirements^1^. This energy – mainly glucose – is delivered to the brain via the cerebral microvasculature and traverses the blood-brain barrier into the parenchyma through the brain endothelial glucose transporter1 (GLUT1) protein^2^. Disrupting the supply of glucose to the brain has dire consequences for this organ, depriving cerebral neurons of their principal energy source and triggering profound dysfunction of neuronal circuits^3^. The “neuroglycopenia” that provokes brain dysfunction is observed in a variety of conditions including Alzheimer’s disease (AD) and traumatic brain encephalopathies (TBEs)^3–5^. However, it is best illustrated and perhaps most conveniently investigated in the monogenic, neurodevelopmental brain energy failure condition, Glut1 deficiency syndrome (GLUT1DS)^6^.

With an incidence as high as 1:24,000 newborns, GLUT1 deficiency syndrome (GLUT1DS) is caused by *de novo* heterozygous mutations or haploinsufficiency of the *SLC2A1 (GLUT1)* gene and, correspondingly, low levels of the GLUT1 protein^7–9^. Complete ablation of the transporter has never been observed in humans and likely results in death *in utero*. Indeed, mutant mice with homozygous mutations in the *Slc2a1 (Glut1)* gene are rarely detected beyond embryonic day 12, whereas animals with only one intact copy of the gene model human GLUT1DS^10^. In humans, the condition becomes apparent in infancy and is characterized by a complex phenotype involving intractable epileptic seizures, a movement disorder that combines elements of ataxia, dystonia and spasticity, and neurodevelopmental delay^11,12^. Exercise-induced dyskinesias and hemolytic anemias complicate GLUT1DS symptoms^13,14^. Yet, all patients exhibit abnormally low levels (<40mg/dL in 90% of patients) of cerebrospinal fluid (CSF) glucose – evidence of neuroglycopenia – and CSF lactate levels in the low to low-normal range (<9mg/dL)^6^.

GLUT1DS was initially described in 1991 and its underlying cause reported in 1998 (refs. 7, 8). Yet, patients afflicted with the condition have few treatment options^15^. The most commonly prescribed therapy is a ketogenic diet. Such diets mitigate seizures in young patients but are less effective in preventing the movement disorders characteristic of GLUT1DS^6,16^. One likely explanation for the modest effects of ketogenic diets is that while they supply the brain with an alternate energy source, ketone bodies, which are fed into the tricarboxylic acid (TCA) cycle to generate energy, they do not address the underlying cause – low brain GLUT1 and deficient cerebral glucose – of the condition and fail to replenish glycolytic intermediates to the cell. One such intermediate, lactate, is reported to be especially important for neuronal function^17^. A straightforward solution to this predicament is to restore GLUT1 levels to the GLUT1 deficient brain. In this study, we have done just that in a mouse model of the disease by transgenically expressing a genomic fragment harboring the human *GLUT1* gene. We show that in mutants expressing the transgene, GLUT1 deficiency disease burden is significantly reduced. Motor performance in such mutants improved, CSF glucose and CSF lactate levels rose, seizure activity subsided and levels of GLUT1 in brain tissue were enhanced. Interestingly, the presence of the transgene in GLUT1DS mutants resulted not only in expression of the human *GLUT1* gene but also in induction of the intact murine *Slc2a1* allele in the animals. One explanation for the latter observation is the effect of a regulatory RNA in the transgene bearing the human *GLUT1* locus. Overall, our results attest to the mitigating effects of restoring GLUT1 to the GLUT1 deficient brain. The transgenic mouse lines described here also constitute a useful pre-clinical tool to test agents that modulate *human GLUT1* expression in an intact mammalian organism.

## Results

### A transgene bearing the human GLUT1 gene locus reduces disease burden in GLUT1DS model mice

An intuitively appealing treatment for GLUT1DS, and one that addresses the root cause of the condition is to raise GLUT1 and thus brain glucose to therapeutic levels. To test this idea, we engineered mice to express a human *GLUT1* transgene and sought to examine the effects of expressing the transgene in a well-established mouse model of GLUT1DS^10^. In anticipation of using the transgenic mice in future studies designed to test agents that modulate human *GLUT1* expression in the intact organism via effects on gene regulatory elements, we opted to generate mice bearing the *entire* human *GLUT1* gene locus rather than just a *GLUT1* cDNA transgene. Accordingly, a bacterial artificial clone (BAC) harboring human *GLUT1* was identified and a ∼79kb fragment containing all 10 exons of the gene and ∼43.5kb of upstream sequence isolated to generate the transgenic mice (Fig. 1A, B). Of note, the region of the transgene located upstream to *GLUT1* contains genomic sequence that codes for a natural antisense transcript (NAT), *SLC2A1-DT*, reported to regulate *GLUT1* expression^18^. Microinjection of the 79kb transgene bearing the human *GLUT1* locus into fertilized mouse oocytes resulted in several lines. However, only two – lines 2829 and 2857 – were found to not only carry the intact transgene (Fig. 1C) but also transmit it through the germline, and these were employed for most of our analysis.

**Figure 1.**
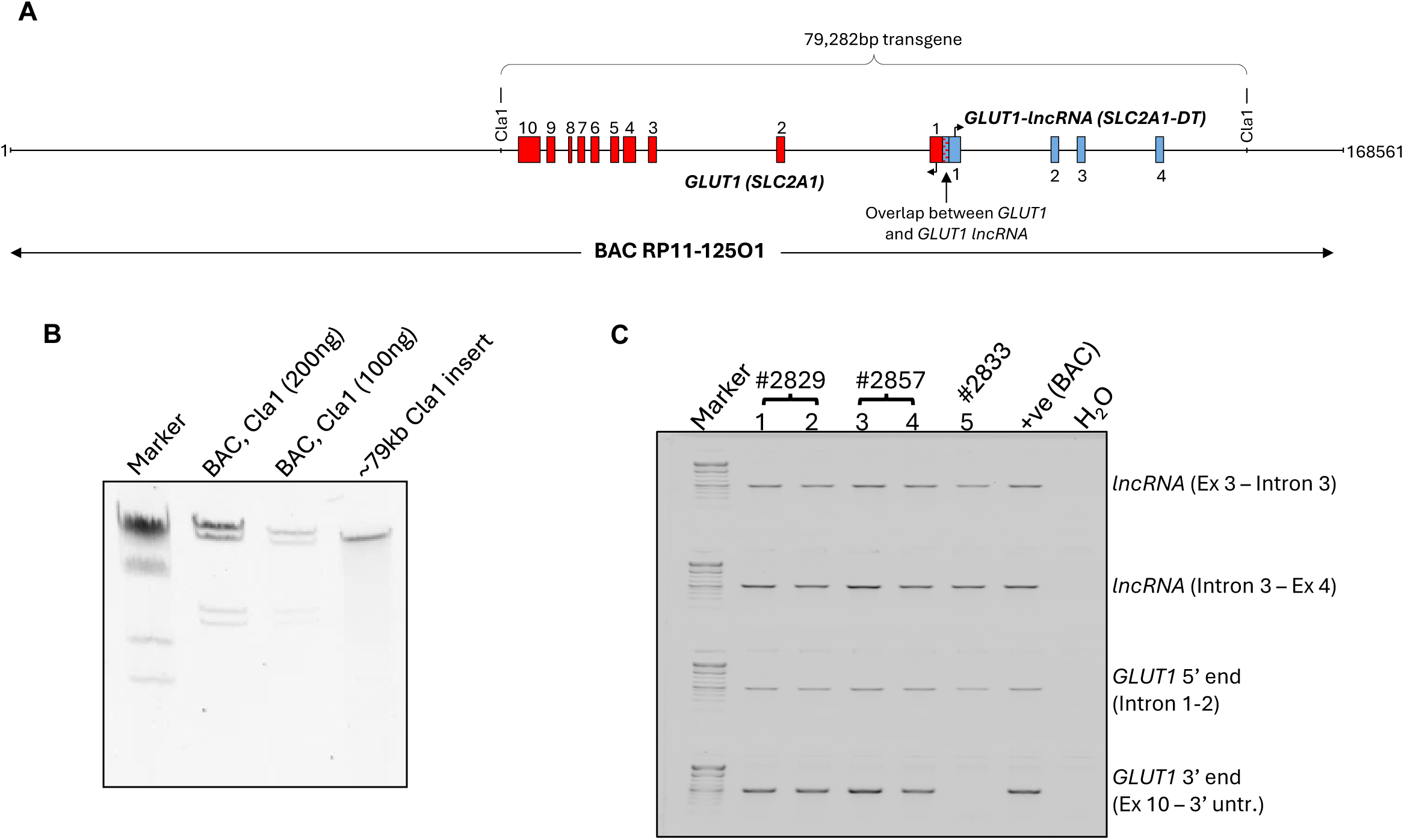
Generation of transgenic mice harboring the genomic human *GLUT1* gene locus. **(A)** Schematic illustrating BAC RP11-125O1 and the Cla1 fragment containing human *GLUT1* and the *SLC2A1-DT* lncRNA used to create transgenic mice. **(B)** Agarose gel showing BAC RP11-125O1 digested with Cla1 and an aliquot of the purified 79kb transgene that was microinjected into fertilized oocytes to derive the transgenic mice. Note: Marker – Lambda DNA digested with HindIII. **(C)** Gel depicting the results of genotyping experiments to determine if the various lines that were produced harbored intact transgenes. Unlike lines 2829 and 2857, line 2833 lacks the 3’ end of the *GLUT1* gene. Note: Marker – 1kb Plus DNA ladder (Thermo Scientific Inc.).

Having identified transgenic lines that harbored the intact 79kb human *GLUT1* locus, we sought to determine if the presence of the transgene could mitigate the severity of the GLUT1DS phenotype. Accordingly, the transgenes in each line were introduced into a GLUT1 deficient background by breeding mice from these lines to a well-established mouse model of GLUT1DS^10^. To ensure that strain heterogeneity did not confound our analysis, each of the lines was first backcrossed over 5 generations with 129 S6/SvEvTac strain mice; GLUT1DS model mice have been maintained on this strain background historically. Progeny GLUT1DS model mice harboring the transgenes from lines 2829 *(2829^Tg^;Glut1^+/-^)* or 2857 *(2857^Tg^;Glut1^+/-^)* were then assessed along with wild-type (WT – *Glut1^+/+^*) and *Glut1^+/-^* controls using a battery of assays to determine phenotypic outcomes. We began by examining motor performance in the four mouse cohorts. In a rotarod test, *Glut1^+/-^* mutants, expectedly, performed significantly worse than WT littermates (Fig. 2A). In contrast, *2829^Tg^;Glut1^+/-^* and *2857^Tg^;Glut1^+/-^* mice exhibited improved performance compared to *Glut1^+/-^* mutants. Moreover, at the two time-points at which animals were tested, the performance of *2829^Tg^;Glut1^+/-^* and *2857^Tg^;Glut1^+/-^* mice was statistically equivalent to that of WT littermates.

**Figure 2.**
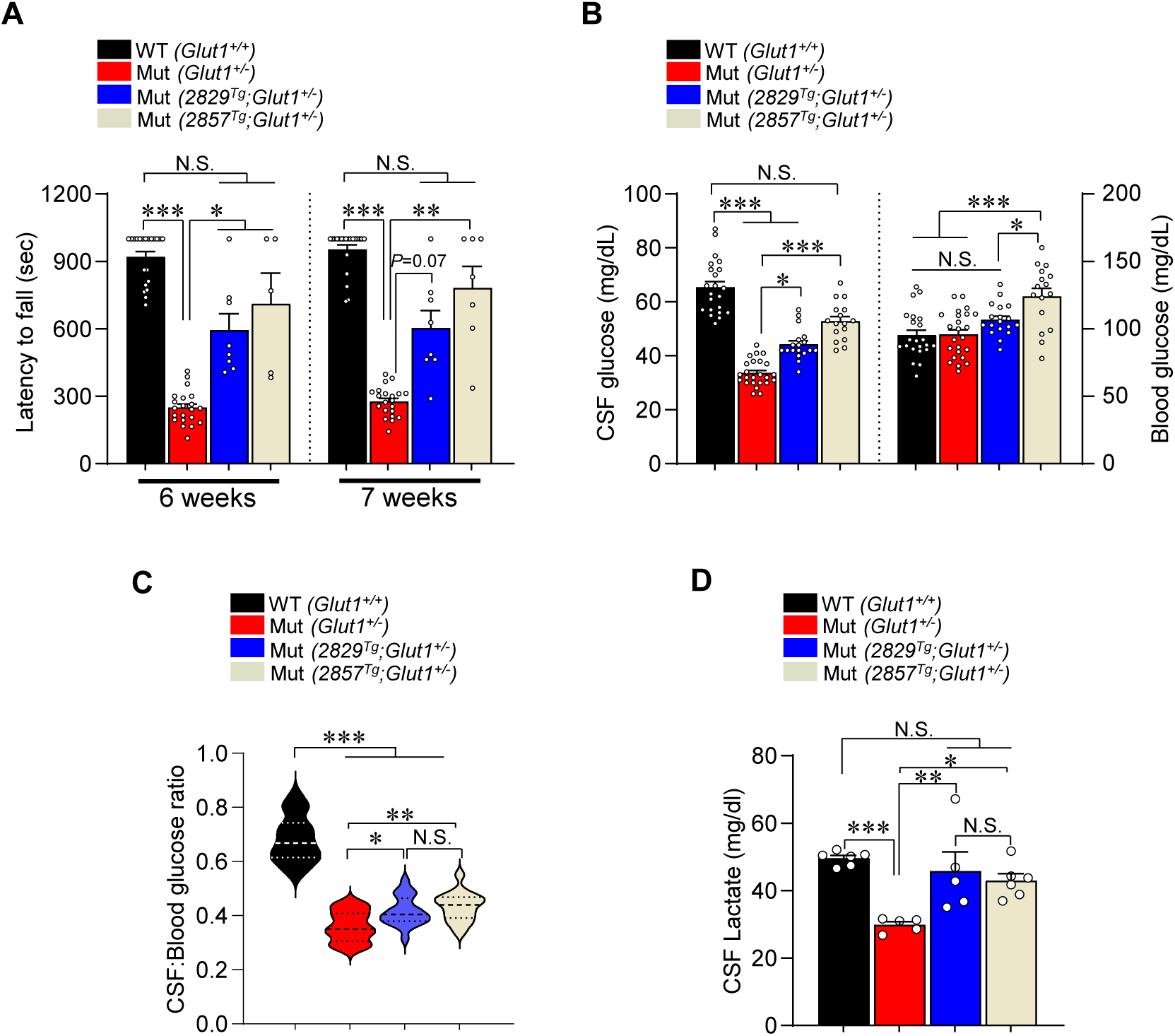
Mitigation of the GLUT1DS phenotype in mutant mice bearing the human *GLUT1* gene locus. **(A)** Quantified outcome of mouse motor performance on a rotarod. Note: *, **, ***, *P* < 0.05, *P* < 0.01 and *P* < 0.001, Kruskal-Wallis test, n = 5 – 21 subjects tested. **(B)** Quantification of CSF and blood glucose levels in controls and GLUT1DS mutants transgenic for the human *GLUT1* gene locus. Note: *, ***, *P* < 0.05, *P* < 0.001; one-way ANOVA and Kruskal-Wallis tests to evaluate means for CSF and blood values respectively, n ≥ 16 mice of each cohort. **(C)** Graph depicting CSF:blood glucose ratios in controls and GLUT1DS mutants bearing the human *GLUT1*-containing transgene. Note: *, **, ***, *P* < 0.05, *P* < 0.01 and *P* < 0.001, one-way ANOVA, n ≥ 16 mice of each genotype. **(D)** Graphical representation of CSF lactate values in controls and GLUT1DS mutants transgenic for the human *GLUT1* gene locus. Note: *, **, ***, *P* < 0.05, *P* < 0.01 and *P* < 0.001, one-way ANOVA, n ≥ 5 mice of each genotype.

Hypoglycorrhachia (low CSF glucose) is a signature feature of human GLUT1DS and is a characteristic we have previously documented in model mice^18–20^. We were therefore not surprised that CSF glucose levels were significantly lower in *Glut^+/-^* mutants than they were in WT *(Glut1^+/+^)* controls (Fig. 2B). Transgenic expression of human *GLUT1* raised CSF glucose significantly in *Glut1^+/-^* mutants, although concentrations remained lower than they were in WT controls (Fig. 2B). Hypoglycorrhachia in GLUT1DS is observed in the context of normoglycemia^6^. A significant increase in blood glucose levels in *2857^Tg^;Glut1^+/-^* mutants was therefore unexpected. Still, the increase in CSF glucose in mutants expressing human *GLUT1* was reflected in significant increases in CSF:blood glucose ratios in each of the lines (Fig. 2C). Hypoglycorrhachia in GLUT1DS is frequently accompanied by hypolactorrhachia (low CSF lactate)^21^. Accordingly, we also assessed lactate concentrations in CSF of our various mouse cohorts. As expected, *Glut1^+/-^* mice exhibited significant hypolactorrhachia (Fig. 2D) relative to WT *(Glut1^+/+^)* littermates. In contrast, we found that CSF lactate levels had been normalized in mutants expressing the 2829 and 2857 human *GLUT1* transgenes. This suggests that notwithstanding somewhat lower levels of CSF glucose in *2829^Tg^;Glut1^+/-^* and *2857^Tg^;Glut1^+/-^* mice – relative to WT *(Glut1^+/+^)* controls, the increase in CSF glucose, compared to non-transgenic *Glut1^+/-^* mutants, is sufficient to normalize lactate concentrations.

Infantile-onset epileptic seizures are often the first recognized signs of brain dysfunction in patients with GLUT1DS. In model mice, seizures manifest as abnormal spike-wave discharges (SWDs) on electroencephalograms (EEGs)^10,19^. Expectedly, we found that *Glut1^+/-^* mutants had significantly greater numbers of SWDs than WT *(Glut1^+/+^)* controls (Fig. 3A). Expressing the human *GLUT1* locus in *Glut1^+/-^* mutants reduced SWDs relative to these abnormal discharges in mutants devoid of the human transgene. This largely normalized EEG activity (Figs. 3B, C), suggesting that the human *GLUT1* transgene does indeed ameliorate disease. In aggregate, the outcome of our various tests constitutes compelling evidence that augmenting *GLUT1* by expressing the human *GLUT1* gene in a GLUT1 deficient background is salutary and reduces the burden of disease as assessed in GLUT1DS model mice.

**Figure 3.**
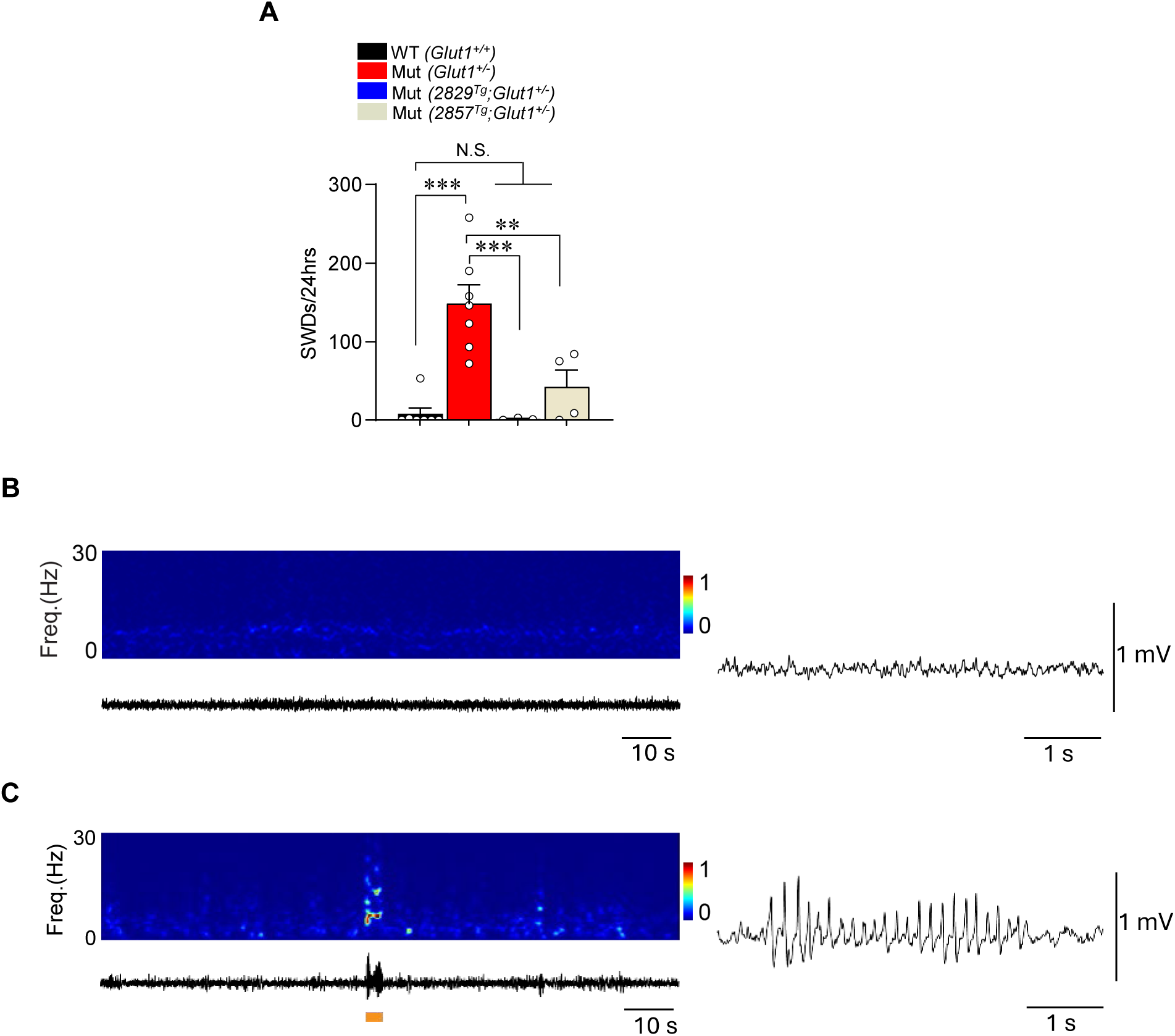
Seizures are suppressed in GLUT1DS mice expressing the human *GLUT1* gene. **(A)** Quantification of spike-wave discharges (SWDs) in controls and GLUT1DS mutants transgenic for the human *GLUT1* gene locus. Note: *, **, ***, *P* < 0.05, *P* < 0.01 and *P* < 0.001, one-way ANOVA, n = 3 – 6 mice of each genotype. Representative EEG spectrograms (at 0-30Hz) and corresponding traces below and to the right of them depicting **(B)** a relatively normal EEG wave pattern obtained from a *2829^Tg^;Glut1^+/-^* subject and **(C)** one from a *2857^Tg^;Glut1^+/-^* mouse exhibiting a SWD – highlighted by orange box – sometimes seen in this cohort of mice.

### The human GLUT1 transgene raises GLUT1 levels in part by stimulating Slc2a1 activity

To ascertain how disease burden in *2829^Tg^;Glut1^+/-^* and *2857^Tg^;Glut1^+/-^* mutants is lowered, we examined the expression of GLUT1 in these two lines of mice. We began by inquiring if human *SLC2A1 (GLUT1)* is expressed in the mice. RT-PCR demonstrated that the human gene is indeed expressed from the 79kb transgenic fragment in brain tissue of subjects from lines 2829 and 2857 but not in mice originating from a third founder line, 2833, determined to be truncated for the transgene and devoid of *SLC2A1* exons 3 – 10 (Fig. 4A). Quantitative RT-PCR confirmed expression of human *GLUT1* in lines 2829 and 2857 and furthermore revealed that the latter expresses the transporter more abundantly (Fig. 4B). Nevertheless, relative to murine *Slc2a1 (Glut1)* transcript levels, the human *GLUT1* gene in the two lines was expressed at low levels – 17% and 31% of murine *Slc2a1* in lines 2829 and 2857 respectively (Fig. 4C). RT-PCR also demonstrated that the *SLC2A1-DT* lncRNA embedded within the 79kb transgene is expressed in the two lines (Fig. 4D). Moreover, line 2857 was found to express modestly greater levels of the lncRNA than line 2829 (Fig. 4E). During the course of these assessments, we also examined murine *Slc2a1 (Glut1)* levels in the two transgenic lines and WT *(Glut1^+/+^)* controls. Interestingly, we found that murine *Glut1* RNA was significantly higher in lines 2829 and 2857 than it was in *Glut1^+/+^* controls (Fig. 5A). While this outcome was initially puzzling, we showed in separate studies that stimulation of murine *Slc2a1* in mice bearing the human *GLUT1* locus is a likely effect of the *SLC2A1-DT* lncRNA in the transgene^18^. Indeed, in brain tissue of *Glut1^+/+^* mice from line 2833, which harbors the intact lncRNA, we found a modest but statistically significant increase in Glut1 protein (Fig. S1A, B). This effect of the lncRNA was confirmed by quantitative RT-PCR analysis of *Glut1* transcripts in GLUT1DS mutant mice with or without the 2833 transgene (Fig. S1C). As expected, *2833^Tg^;Glut1^+/-^* mice expressed robust levels of the lncRNA, whereas the transcript was undetectable in GLUT1DS mutants devoid of the transgene (Fig. S1D). Consistent with a stimulatory effect of the lncRNA on murine *Slc2a1*, hypoglycorrhachia in GLUT1DS model mice harboring the 2833 transgene was mitigated relative to this condition in mutants absent the transgene (Figs. S1D, E). Collectively, the outcomes observed in the various transgenic lines suggest that higher Glut1 levels and reduced GLUT1 deficiency disease burden in mutants from lines 2829 and 2857 result not only from expression of human *GLUT1* in the mice but also from induction of murine *Slc2a1* by the *SLC2A1-DT* lncRNA contained in the 79kb transgene used to generate our transgenic animals.

**Figure 4.**
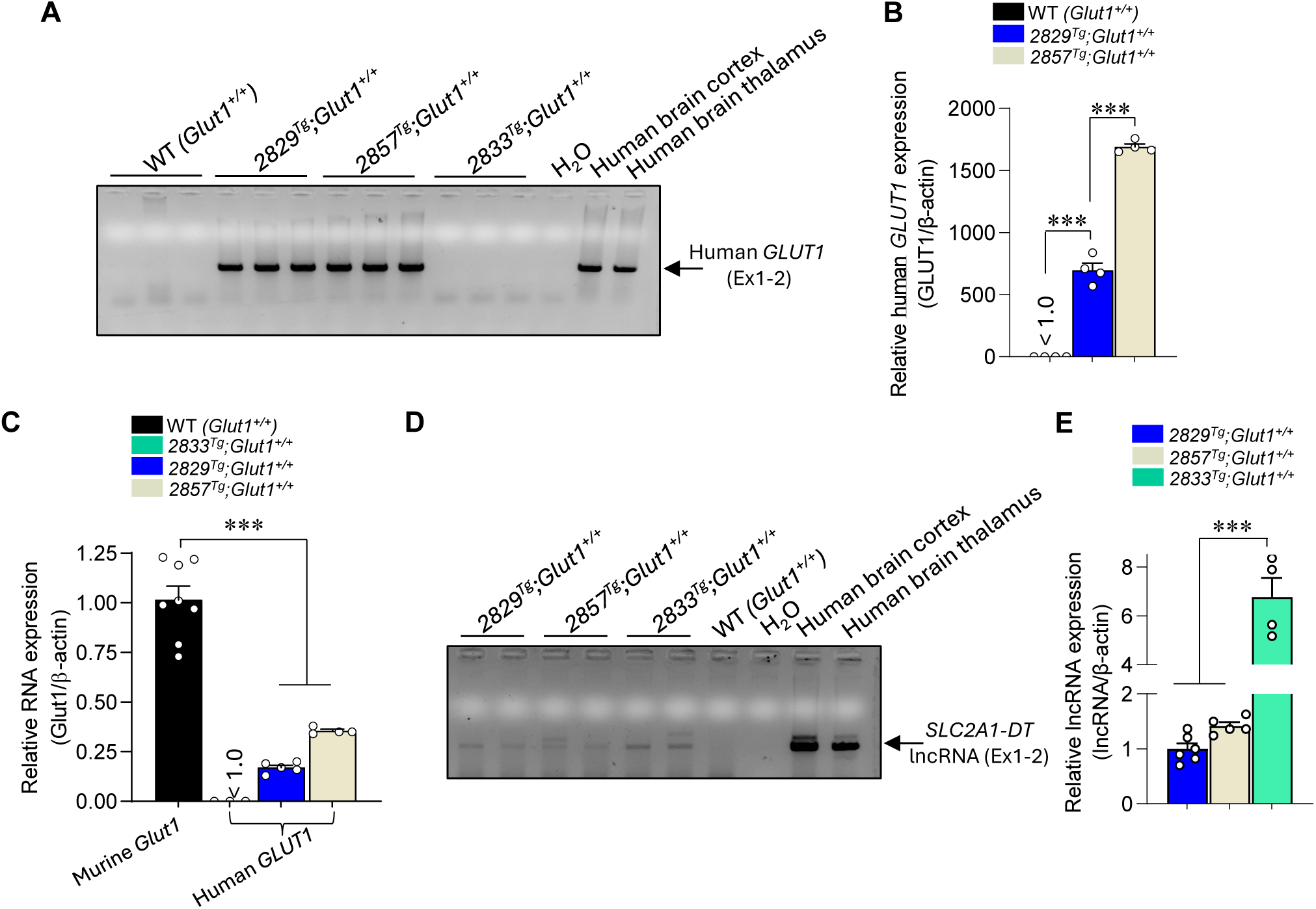
Expression of *GLUT1* and the *SLC2A1-DT* lncRNA from the 79kb transgene. **(A)** Agarose gel showing bands representing RT-PCR products – from brain tissue – originating from the human *GLUT1* gene in the 79kb transgene. Note that only mice from lines 2829 and 2857, carrying the intact transgene, express *GLUT1*. **(B)** Quantified levels of the human *GLUT1* gene in 2829 and 2857 transgenic mice. Note: ***, *P* < 0.001, one-way ANOVA, n = 4 mice of each genotype. **(C)** Graph depicting levels of human *GLUT1* expression relative to those of its murine *Slc2a1 (Glut1)* counterpart in brain tissue of relevant mice. Note: ***, *P* < 0.001, one-way ANOVA, n = 3 – 8 mice of each genotype. **(D)** Agarose gel showing bands representing RT-PCR products – in brain tissue – from the human *SLC2A1-DT* lncRNA-expressing element in the 79kb transgene. In this instance, all three transgenic lines expressed the lncRNA. **(E)** Quantified levels of the human lncRNA in the three lines of transgenic mice. Note: ***, *P* < 0.001, one-way ANOVA, n ≥ 4 mice of each genotype.

**Figure 5.**
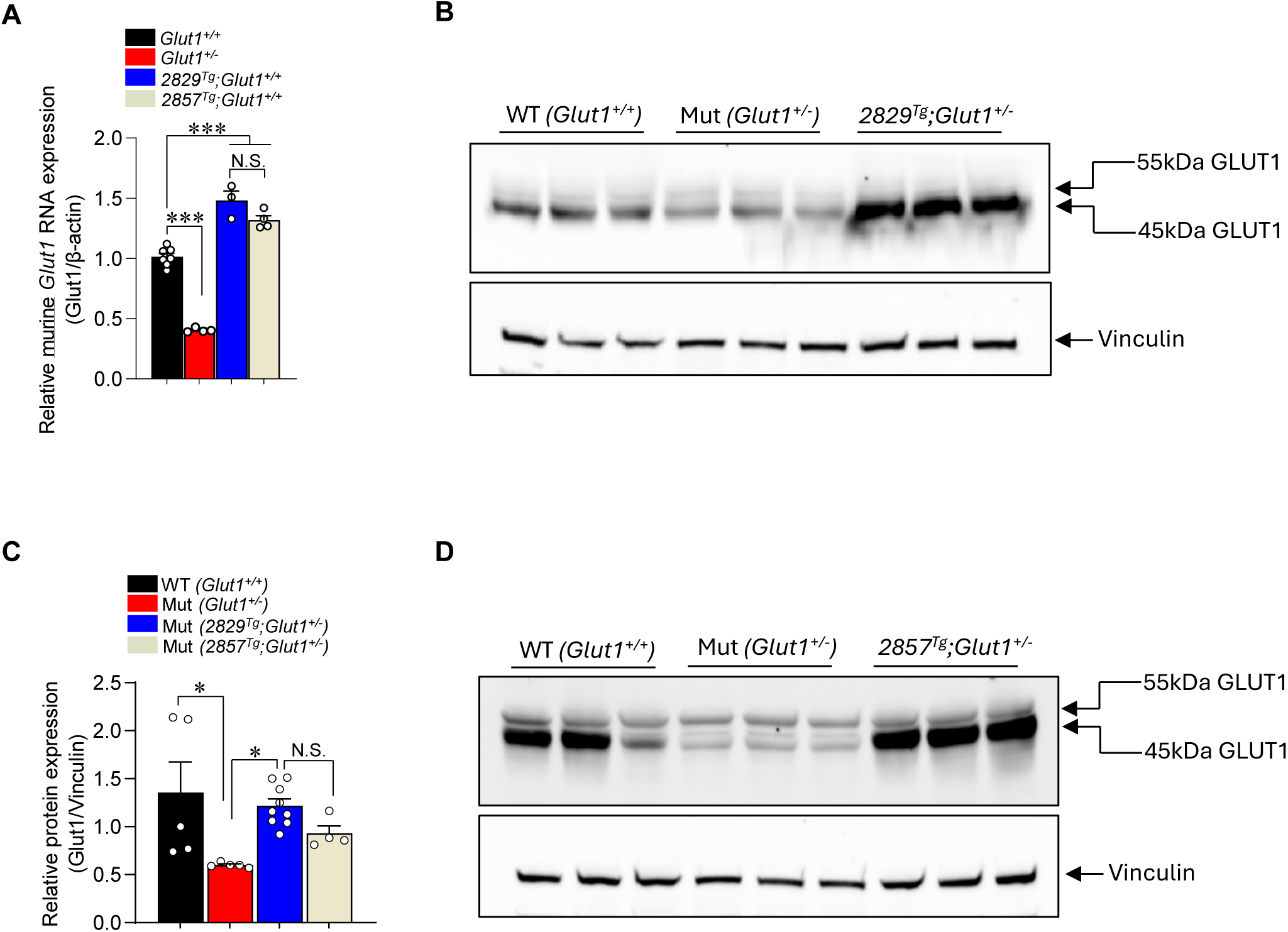
Transgenic expression of the human *GLUT1* locus raises murine *Slc2a1* and total GLUT1 levels. **(A)** Quantified levels of murine *Slc2a1 (Glut1)* transcripts in brain tissue of WT and transgenic mice. Note: ***, *P* < 0.001, one-way ANOVA, n ≥ 3 mice of each genotype. **(B)** Representative western blot of total GLUT1 protein in brain tissue of controls, and mutants expressing the human *GLUT1* locus in line 2829. Note: 55kDa band – endothelial GLUT1; 45kDa band – astrocytic GLUT1. **(C)** Quantified levels of total GLUT1 protein in controls and mutants expressing the human *GLUT1* locus in lines 2829 and 2857. Note: *, *P* < 0.05, one-way ANOVA, n ≥ 4 mice of each genotype. **(D)** Representative western blot of total GLUT1 protein in brain tissue of controls, and mutants expressing the human *GLUT1* locus in line 2857. Note: 55kDa band – endothelial GLUT1; 45kDa band – astrocytic GLUT1.

Considering mitigation of the GLUT1DS phenotype in *2829^Tg^;Glut1^+/-^* and *2857^Tg^;Glut1^+/-^* mutants and expression not only of human *GLUT1* but also stimulation of murine *Slc2a1 (Glut1)* in the mice, we predicted an increase in total GLUT1 protein in these animals. True to expectation, western blot analysis of brain tissue from *2829^Tg^;Glut1^+/-^* mutants revealed that these animals did indeed express more transporter than GLUT1DS mice that did not harbor the transgene (Fig. 5B, C). Consistent with prior studies, a significant difference in protein levels was also observed between WT *(Glut1^+/+^)* controls and *Glut1^+/-^* mutants. Similar patterns of GLUT1 protein expression were observed when we compared *2857^Tg^;Glut1^+/-^* mutants with control mice (Fig. 5C, D). In aggregate, these results serve as important evidence that restoring GLUT1 to GLUT1DS subjects offers protection against disease.

### Expression of the human GLUT1 locus in lines 2829 and 2857 is insufficient to prevent Glut1^-/-^ lethality

Complete ablation of Glut1 *(Glut1^-/-^)* is embryonically lethal^10^. Considering the ability of the human *GLUT1* locus in lines 2829 and 2857 to mitigate GLUT1DS disease severity, we inquired if expression of the locus might rescue the embryonic lethality resulting from homozygous loss of the murine *Glut1* gene. Accordingly, we first bred *2829^Tg/Tg^;Glut1^+/-^* mice with each other to ascertain if the 2829 transgene on a murine *Glut1* null background *(2829^Tg/Tg^;Glut1^-/-^)* might produce viable animals; breeders were made homozygous for the transgene to reduce the various possible genotypes in progeny mice. Of 86 pups generated from such crosses, none were found to be null for murine *Glut1* (Fig. 6A). This outcome deviated significantly from the expected frequency of 25% from the experimental crosses (*P* < 0.0001, df=2, χ^2^=28.69) suggesting that levels of transporter from the human *GLUT1* transgene in line 2829 are insufficient to prevent embryonic lethality, notwithstanding mitigation of disease on the *Glut1* heterozygous background. We assessed the 2857 transgene similarly, except that the breeders employed to generate pups were heterozygous for the transgene *(2857^Tg/-^;Glut1^+/-^)*. While this increased the number of possible genotypes, akin to our analysis of line 2829, we failed to generate *Glut1^-/-^* mutants with or without the 2857 transgene (Fig. 6B). Of 79 viable pups obtained, none were found to be null for the murine *Glut1* gene. This too deviated significantly from an expected frequency of ∼20% (*P* < 0.0001, df=7, χ^2^=45.49), suggesting that the 2857 transgene also likely expresses insufficient GLUT1 in the absence of a normal *Slc2a1* copy to ensure live births.

**Figure 6.**
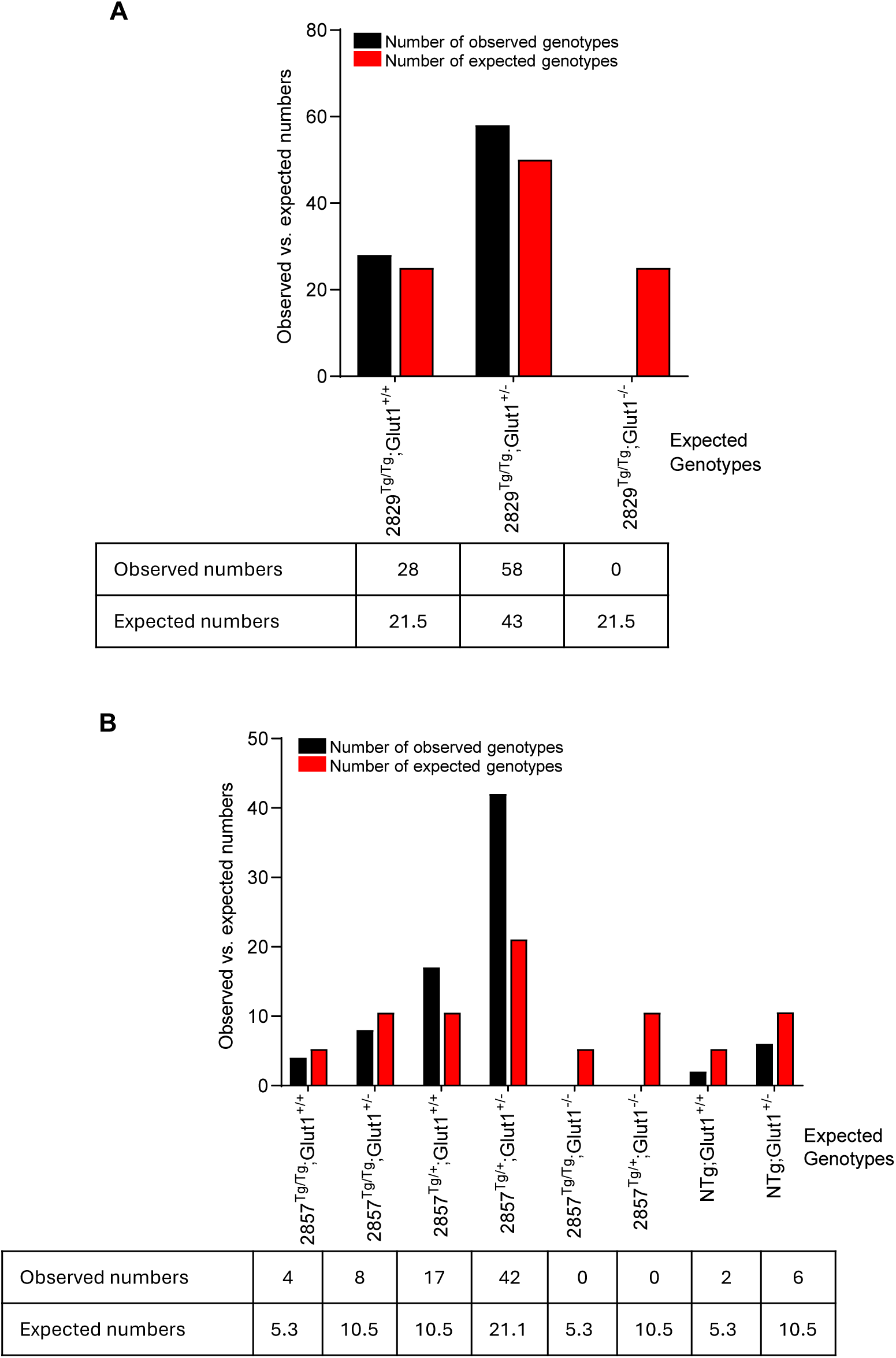
Transgenic expression of the human *GLUT1* locus in lines 2829 and 2857 does not compensate for absence of murine *Glut1.* **(A)** Graphical representation of observed and expected genotypes resulting from crosses between *Glut1* haploinsufficient mice homozygous for the 2829 transgene. Mouse numbers are tabulated below the graph. **(B)** Graphical representation of observed and expected genotypes resulting from crosses between *Glut1* haploinsufficient animals heterozygous for the 2857 transgene. Mouse numbers are tabulated below the graph. Note: The numbers in the two tables correspond to the genotypes on the x-axes of the graphs above them.

While attempting to determine if the expression of the human *GLUT1* locus in transgenic line 2857 rescued *Glut1^-/-^* embryonic lethality, we noticed that 5-month-old progeny homozygous for the 2857 transgene, irrespective of whether they harbored one or two WT murine *Glut1* alleles, were significantly smaller than age-matched littermates heterozygous for the transgene [2857 homozygotes: 18.95g ± 0.84g versus 2857 heterozygotes: 31.86g ±1.02g, *P* < 0.001, *t* test, n ≥ 7 male mice of each cohort]. In contrast, mice homozygous for the 2829 transgene were visually indistinguishable from healthy and WT controls. Suspecting that the dwarf phenotype of the 2857 homozygotes stemmed from an unintended transgene insertion effect that was line-specific and unmasked only in the homozygous state, we sought to map the 2857 insertion site. To do so, Vectorette PCR, which combines transgene-specific primers with adaptor primers, was employed^22,23^. We found that the 2857 transgene had inserted into intron 7 of the Pappalysin *(Pappa)* gene located on Chr. 4 (Fig. S2A). *Pappa* knockout mice are viable and fertile but consistent with our own observations, reported to be significantly smaller than WT littermates^24^. To determine if insertion of our *GLUT1* transgene had indeed rendered the *Pappa* gene in line 2857 inactive, we carried out RT-PCR to amplify *Pappa* transcripts in *Glut1^+/-^* mutants homozygous, heterozygous or devoid of the 2857 transgene. This analysis revealed that while *Pappa* transcripts were apparent in *Glut1^+/-^* and *2857^Tg/-^;Glut1^+/-^* mice, they were undetectable in *Glut1^+/-^* mutants homozygous *(2857^Tg/Tg^;Glut1^+/-^)* for the transgene (Fig. S2B). Consistent with these findings, quantitative RT-PCR demonstrated that mice heterozygous for the 2857 transgene *(2857^Tg/-^;Glut1^+/-^)* expressed only half as much *Pappa* RNA as did individuals WT for this gene, while animals homozygous for the transgene expressed even less (Fig. S2C). These results suggest that *Pappa* is indeed rendered non-functional by the *GLUT1* transgene in line 2857. While such an outcome was clearly unintended, it not only explains the smaller size of 2857 homozygotes but is also consistent with the somewhat hyperglycemic state of animals harboring this transgene – also see Fig. 2B); *Pappa* knockout reportedly raises blood glucose levels significantly via effects on adipose tissue, which plays a central role in the regulation of insulin sensitivity^25^.

## Discussion

Notwithstanding its initial description more than three decades ago, GLUT1DS remains a disease with an unmet medical need^26^. Moreover, given the biology of the disorder, which involves reduced nutrient delivery to the brain, GLUT1DS also serves as a useful paradigm to investigate the larger family of cerebral energy failure syndromes. Here, we describe work whose twin goals are to better understand neuroglycopenia and, more immediately, to treat the root cause of it in the neurodevelopmental condition GLUT1DS. We expect outcomes reported here to become relevant to several CNS energy failure syndromes including retinitis pigmentosa and AD^4,5,27^. Four noteworthy findings emerge from our study. First, we provide evidence that expression of the human *GLUT1* gene – from a transgene in this instance – augments transporter activity sufficiently in a mouse model of GLUT1DS to mitigate disease severity. This sets the stage for gene-replacement type studies that employ the *human GLUT1* gene to treat GLUT1DS and other neuroglycopenic conditions stemming from disruption of the transporter. A second important outcome of the work described here derives from observations of the effects of the transgenes bearing the human *GLUT1* locus on overall *GLUT1* expression. We documented an intriguing effect of our transgenes on *murine Slc2a1 (Glut1)* expression. An analysis of one of our transgenic lines harboring a truncated *GLUT1* gene but intact *SLC2A1-DT* natural antisense transcript suggests that it is the lncRNA that stimulates *Glut1* activity. This is consistent with a recent report from our laboratory^18^ and provides additional insight into the intricate regulation of *GLUT1* expression. A third notable outcome, albeit one that is serendipitous, involves observations associated with transgenic line 2857. Unexpectedly, the transgene in this line caused a mild hyperglycemic state. While such a condition may be a confounding factor in our analysis, it also raises the prospect of augmenting brain glucose in GLUT1DS in new and unorthodox ways. Finally, our work has resulted in novel transgenic lines that will be useful not only to test, validate and optimize agents designed to alter human *GLUT1* for the treatment of GLUT1DS but also to study how this gene is regulated in an established mammalian model organism.

The most common clinical treatment for GLUT1DS is the ketogenic diet^6^. While the diet suppresses seizures associated with GLUT1 deficiency, its effects on other characteristic features of the disease, notably movement defects and dysarthria are modest at best. This likely derives from several related factors, the most salient of which is that the brain prefers glucose, whereas the ketogenic diet substitutes this nutrient with an imperfect energy source, ketone bodies^28,29^. Ketone bodies function as anaplerotic agents by serving as a source of acetyl CoA for the TCA cycle. However, they have little effect in replenishing glycolytic intermediates, which likely play an important role in brain energetics. Moreover, entry of ketone bodies into the brain declines upon weaning when the switch to glucose as a preferred brain energy source is paralleled by a downregulation of MCT1 (monocarboxylate transporter 1), the conduit through which the ketones traverse the blood-brain barrier^30^. Therapeutic strategies that restore GLUT1 to the GLUT1 deficient brain address the drawbacks of the ketogenic diet. Indeed, we have previously demonstrated that raising *murine Glut1* in GLUT1DS model mice mitigates disease^19^. Here we show that human *GLUT1*, the more intuitive choice for a clinical gene-replacement therapeutic strategy, has a similar effect in transgenic mice.

Using two independent lines of transgenic mice harboring a ∼79kb human *GLUT1* gene locus, we showed that brain GLUT1 levels could be raised in a mouse model of GLUT1DS. This had the effect of improving motor performance, combating hypoglycorrhachia, normalizing CSF lactate levels and lowering seizure activity. Failure to restore mutant mice to full health is a likely consequence of sub-optimal *GLUT1* expression from the transgenes, most consequentially in critical cellular sites of action of the transporter. While such an outcome is a recognized caveat of expressing genes transgenically, it is usually remedied with the generation of additional lines representing an allelic series. Still, our results using lines 2829 and 2857 inform minimum levels of GLUT1 needed to ensure live births. For instance, If *GLUT1* in the former line is truly expressed at ∼17% of WT levels as assessed in this study, and if these levels failed to yield any *2829^Tg/Tg^;Glut1^-/-^* mutants, one might infer that gestational viability requires expression of the transporter at greater than 34% of normal amounts. Given the higher expression of human *GLUT1* in line 2857 (∼60% of WT GLUT1, when homozgous), one would have expected this transgene to rescue *Glut1^-/^*^-^ embryonic lethality. Failure to detect viable *2857^Tg/Tg^;Glut1^-/-^* mutants was therefore surprising but may be explained by the additional disease burden placed on these animals because they are perforce knocked out for the *Pappa* gene. Pertinent to this reasoning is data reporting that the Pappa protein is a critical growth regulatory factor during fetal development^24^.

*Pappa* knockout by the 2857 transgene and consequent hyperglycemia in mice bearing the transgene could confound our conclusions about the suppression of GLUT1DS in *2857^Tg^;Glut1^+/-^* mutants, particularly as it relates to mitigation of the hypoglycorrhachia in these mutants; increased CSF glucose in these animals is a likely combination of higher GLUT1 levels *and* modest hyperglycemia. The relative salutary effect of each is unclear. Yet, it raises the intriguing possibility of using a mild hyperglycemic state to combat neuroglycopenia. Considering how glucose traverses the GLUT1 transporter – by facilitated diffusion, which is subject to the concentration gradient of the nutrient on the luminal versus abluminal side of the brain microvasculature, one might indeed envisage driving glucose entry into the GLUT1 deficient brain by raising blood glucose levels. Proof-of-concept studies to investigate such a mechanism is the focus of future work that takes advantage of independent *Pappa* knockout mice^31^ to examine the effects of the knockout on the GLUT1DS phenotype. The feasibility of raising blood glucose modestly to treat GLUT1DS, particularly in patients intolerant to the ketogenic diet, is bolstered by encouraging results when they are administered diazoxide, a small molecule normally used to suppress hyperinsulinism^32,33^.

A second serendipitous outcome of this study worth highlighting stems from the effect of our various transgenes on murine *Glut1* expression. Stimulation of the murine *Slc2a1* gene by the transgenes may be puzzling but is consistent with recent work from our lab identifying a regulatory lncRNA, *SLC2A1-DT*, in the human *GLUT1* locus that concordantly raises expression from its protein coding cognate^18^. That it is indeed the lncRNA that produces this effect rather than human *GLUT1* itself is supported by our analysis of line 2833. This line, which was found to be truncated for *GLUT1* but retained an intact copy of the lncRNA, was nevertheless able to induce murine *Slc2a1* expression akin to that observed in the lines containing the complete transgene. The *GLUT1*-inducing effects of the human lncRNA combined with its ability to do so in *trans* is noteworthy and adds to the repertoire of molecules engaged in tuning *GLUT1* expression. Ongoing studies seek to understand the mechanism of action of the lncRNA. Notwithstanding our limited understanding of molecular mechanisms associated with the lncRNA and several interesting questions raised by this work, our study has shown that a healthy copy of the human *GLUT1* gene can effectively raise GLUT1 levels to treat neuroglycopenia – as assessed in GLUT1DS model mice. Additionally, the transgenic lines we report here constitute new tools to test agents designed to modulate the human *GLUT1* gene in an intact organism.

## Materials and Methods

### Model mice and human tissues

Haploinsufficient *Glut1^+/-^* model mice were generated at Columbia University and have been described in several studies^10,19,20^. Experimental subjects were generally propagated by breeding mutant male mice with WT females, as *Glut1^+/-^* females are poor mothers. All mice were maintained on a 129 S6/SvEvTac genetic background, and mutant genotypes revealed by PCR analysis. Littermate controls were used in all experiments. Lines 2829^Tg^, 2857^Tg^ and 2833^Tg^ were generated for this study by the Columbia University Genetically Modified Mouse Models Core Facility. Briefly, BAC RP11-125O1 (BACPAC Genomics, Emeryville, CA) containing the human *GLUT1* locus was digested with ClaI (Cat.# R0197S, NEB), and a ∼79kb fragment harboring *SLC2A1* and *SLC2A1-DT* gel extracted and purified. The fragment was then dissolved in microinjection buffer and introduced into fertilized mouse oocytes. Transgenic founders were interbred with *Glut1^+/-^* animals to produce *2829^Tg^;Glut1^+/-^, 2857^Tg^;Glut1^+/-^* and *2833^Tg^;Glut1^+/-^* cohorts; mice used for our experiments were heterozygous for the transgenes. Note that except for weight measurements to assess differences between 2857^Tg/Tg^ and relevant controls, our experiments did not discriminate between males and females. Human tissues were obtained from the New York brain bank.

### Phenotypic evaluations-Rotarod test

For behavioral outcomes, we used GraphPad Prism to determine sample sizes to detect differences of at least two standard deviations with a power of 80% (*P* < 0.05). Motor performance was assessed on an accelerating rotarod (Ugo Basile Inc., Italy) as reported in earlier studies^19,20^. Briefly, we subjected mice to a training period of 5mins. on an accelerating rotarod from 0-40rpm three times a day for four consecutive days. Performance on day 5 was assessed at a setting of 25rpm. The rotarod apparatus was cleaned and dried prior to testing. Latency to fall off the rotating rod was recorded and the experiment terminated if a mouse surpassed 1000s.

### Measurements of glucose and lactate in CSF and/or serum

Prior to assessing CSF glucose/lactate and serum glucose levels, mice were fasted overnight. Blood was collected from the tail vein, while CSF was extracted from the cisterna magna as reported by us in previous studies^19,20^. Glucose concentrations in the CSF and blood were assessed using disposable strips and a Contour Next EZ glucose meter (Bayer Corp., NJ). CSF lactate was measured on an Analox GL5 Analyzer (Analox Instruments, Lunenburg, MA) using a Lactate II reagent kit (GMRD-093, Analox Instruments).

### Quantitative PCR and western blotting

Human and murine *GLUT1* transcripts and *SLC2A1-DT* lncRNA levels were quantified by Q-PCR (see Supplemental Information for primer sequences). RNA was extracted from brain tissue using Trizol (Life Technologies, Carlsbad, CA) according to the manufacturer’s instructions and cDNA synthesized using the RevertAid First Strand cDNA Synthesis Kit (Thermo Scientific). Reactions were performed in a CFX96 Real-Time Systems Q-PCR machine (Bio-Rad Labs) using the SsoAdvanced Universal SYBR Green Supermix (Bio-Rad Labs). Relative quantification of RNA expression was calculated using the ΔΔCT method. GLUT1 protein in the various cohorts of mice was estimated by western blot analysis using standard techniques as previously described^19,20^. Briefly, whole brain tissue was lysed in lysis buffer containing protease inhibitors (Complete Protease Inhibitor Cocktail tablets, Roche, Indianapolis, IA) and 50μg of total protein resolved by gradient SDS-PAGE (Bio-Rad Labs). Blots were initially incubated with GLUT1 rabbit polyclonal (1:5000; Millipore), vinculin monoclonal (1:5000; Abcam) or β-actin monoclonal antibodies. Following several washes to remove non-specifically bound primary antibody, the blots were incubated with HRP-linked goat anti-rabbit (Invitrogen Inc.) or HRP-linked goat anti-mouse (Jackson Immunoresearch) secondary antibodies diluted 1:10,000. Protein bands were visualized on a ChemiDoc Imaging System machine (Bio-Rad Labs, Hercules, CA, USA) using the ECL Detection Kit (Cat.# 1705061, Bio-Rad Labs). Band intensities were assessed using ImageJ software (NIH).

### EEG analysis

Mice were rendered unconscious using a ketamine (100mg/kg), xylanine (10mg/kg) mixture administered intra-peritoneally and then placed on a stereotaxic frame with a closed-loop heating system to maintain body temperature. After asepsis, the skull was exposed. For EEGs, a reference screw was inserted into the skull on top of the cerebellum. EEG recordings were made from two screws on top of the cortex 1mm from midline, 1.5mm anterior to the bregma and 1.5mm posterior to the bregma, respectively. The EEG screws were connected to a PCB board which was soldered with a 5-position pin connector. Implants were secured onto the skull with dental cement (Lang Dental Manufacturing) and animals allowed to recover for a week before being subjected to recordings. EEG recordings were performed for 24-48 hours (light on at 7:00AM and off at 7:00PM) in a behavioral chamber housed inside a sound-attenuating cubicle (Med Associated Inc.). Subjects were habituated in the chamber for at least 4hrs. before initiating recordings. EEG signals were recorded, bandpass filtered at 0.5-500 Hz, and digitized at 1017 Hz with 32-channel amplifiers (TDT, PZ5 and RZ5D or Neuralynx Digital Lynx 4S). To estimate SWDs, FFT of EEG was performed using a 1s sliding window, sequentially shifted by 0.25s increments. Subsequently, “seizure”-power (19-24 Hz) was calculated to extract SWD events based on a threshold of 2-3 standard deviations as described previously^33^. A 19-24 Hz band was selected based on its clear separation from normal brain oscillatory activities, although the primary spectral band of SWDs in mice is around 7Hz, which overlaps with theta oscillations during REM sleep or active periods. Two SWD events were merged into one event if their interval was shorter than 1s. Any SWD event with a duration < 0.5s was removed before conducting the analysis. Algorithm-detected SWD events were confirmed manually by investigators. Mouse behavior was monitored during recordings using an infrared video camera at 30 frames per second.

### Mapping transgene insertion site

We employed vectorette PCR^34,35^ to map the insertion site of the 2857 transgene. For this, genomic DNA from 2857^Tg^ mice was first digested in different reactions with each of the following enzymes – RsaI, HincII, HpaI and ScaI – to create corresponding libraries. Subsequently, an adaptor (vectorette) that exhibits complementarity at each end but not in a central 29bp region was ligated onto blunt ends generated by the various restriction enzymes. Such libraries were subsequently subjected to an initial round of PCR using a transgene specific primer and a vectorette primer (UVP) that binds to sequence in the “bubble” formed within the vectorette. Amplification products were then used as template for a nested PCR using the vectorette primer and a second transgene specific primer. The nested PCR functions to increase PCR specificity and generally resulted in a distinct amplification product. In instances where the nested PCR failed to yield a single distinct PCR product, a double-nested PCR was performed using amplification products from the nested PCR as template. The final PCR products were then sequenced, junction fragments determined and sequence in these fragments corresponding to the mouse genome mapped onto the murine reference genome (GRCM39) to identify the precise insertion site of the transgene.

### Statistics

The choice of parametric or non-parametric statistical tests for study outcomes was determined by first assessing if our data was normally distributed or not. For data that conformed to a Gaussian distribution, the unpaired 2-tailed Student’s *t* test or 1-way ANOVA followed by Tukey’s post-hoc comparison, where indicated, were used to compare means for statistical differences. If the data was not normally distributed, the Mann-Whitney test or Kruskal-Wallis test followed by Dunn’s multiple comparison test were used to compare ranks. Data in the manuscript is reported as mean ± SEM. *P* < 0.05 was considered significant. Statistical analyses were performed with GraphPad Prism v9.0 (GraphPad Software).

### Study approval

The procedures reported here adhere to protocols described in the *Guide for the Care & Use of Lab Animals* (National Academic Press, 2011) and were approved by Columbia University’s IACUC. Mice for this study were randomly selected. Additionally, all subjects were maintained on a 129 S6/SvEvTac genetic background and housed in a controlled environment on a 12hr. light/dark cycle with food and water.

## Data availability

Underlying values associated with the data presented in the study may be requested and will be made available by the lead contact, Umrao R. Monani.

### Acknowledgments

We are grateful to members of the Giblin Laboratories for critically reading this manuscript and to Darryl De Vivo for his insights and unswerving support throughout this investigation. Work described here was funded by the GLUT1 Deficiency Foundation (U.R.M., M.T.), The University of Pennsylvania Orphan Disease Center (U.R.M., M.T.), the Hope for Children Research Foundation (U.R.M.), AFM-France (U.R.M.), the Crofoot family foundation, and NIH (R03 NS128211). Y.P. acknowledges support from the Columbia Precision Medicine Initiative.

## Author contributions

M.T. planned and performed most of the experiments described here. S.T. and Y.P. carried out the EEG analyses. A.Y.K. was involved in animal husbandry and genotyping mice for the study. U.R.M. conceptualized the experiments, directed the project, analyzed and interpreted the data. U.R.M. and M.T. wrote the manuscript with input from all authors.

## Supplemental Information

## Key Reagents

### PCR primers

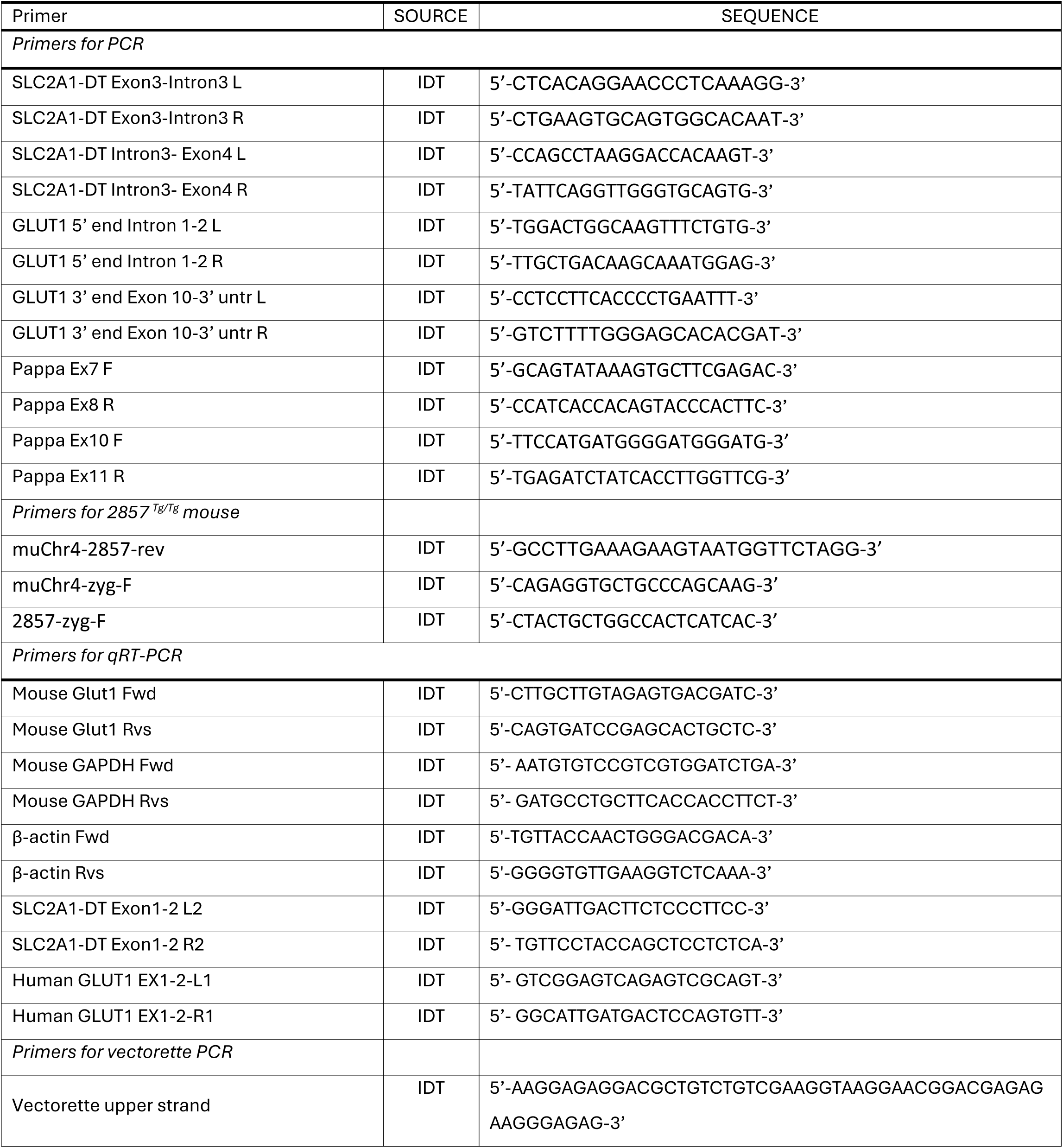
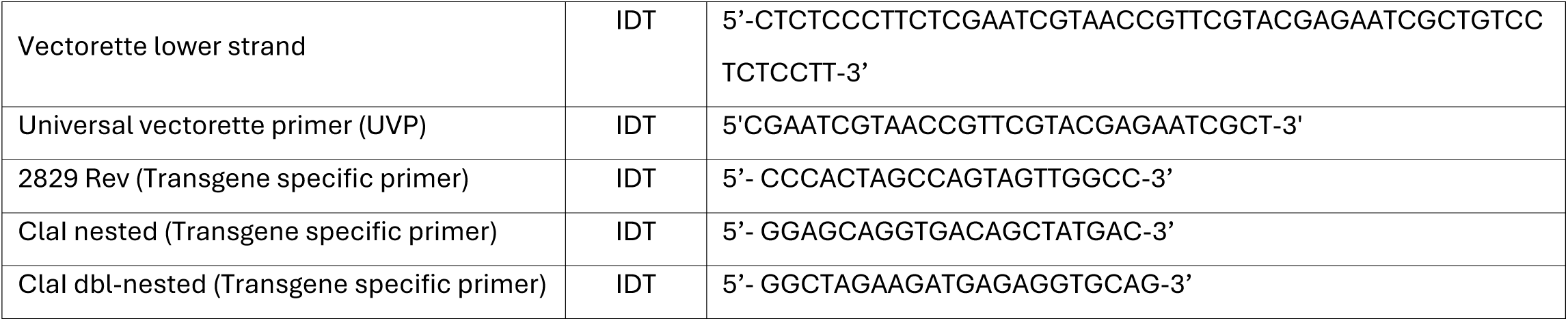

### Other reagents

| REAGENT or RESOURCE | SOURCE | IDENTIFIER |
| --- | --- | --- |
| *Antibodies for Western blotting* |  |  |
| GLUT1 (1:5000) | Millipore | Cat#07-1401 |
| Vinculin (1:5000) | Abcam | Cat#ab129002 |
| β-actin (1:5000) | Sigma | Cat#A5441 |
| Goat anti-rabbit IgG (1:10,000) | Invitrogen | Cat#31460 |
| Goat anti-mouse IgG (1:10,000) | Jackson Immunoresearch | Cat#115-035-003 |
| *Experimental Models: Organisms/Strains* |  |  |
| *Glut1*<sup>+/-</sup> mouse model | Giblin Labs, Columbia University | N/A |
| *SLC2A1-DT*<sup>Tg</sup> transgenic mice (2833<sup>Tg</sup>) | Giblin Labs, Columbia University | N/A |
| *Human SLC2A1* & *SLC2A1-DT*<sup>Tg</sup> transgenic mice (2829<sup>Tg</sup>, 2857<sup>Tg</sup>) | Giblin Labs, Columbia University | N/A |
| *Software and Algorithms* |  |  |
| GraphPad Prism | Graph Pad Software | https://www.graphpad.com/scientificsoftware/prism/ |
| ImageJ | NIH | https://imagej.nih.gov/ij/ |

---

**Supplemental Figure 1.**
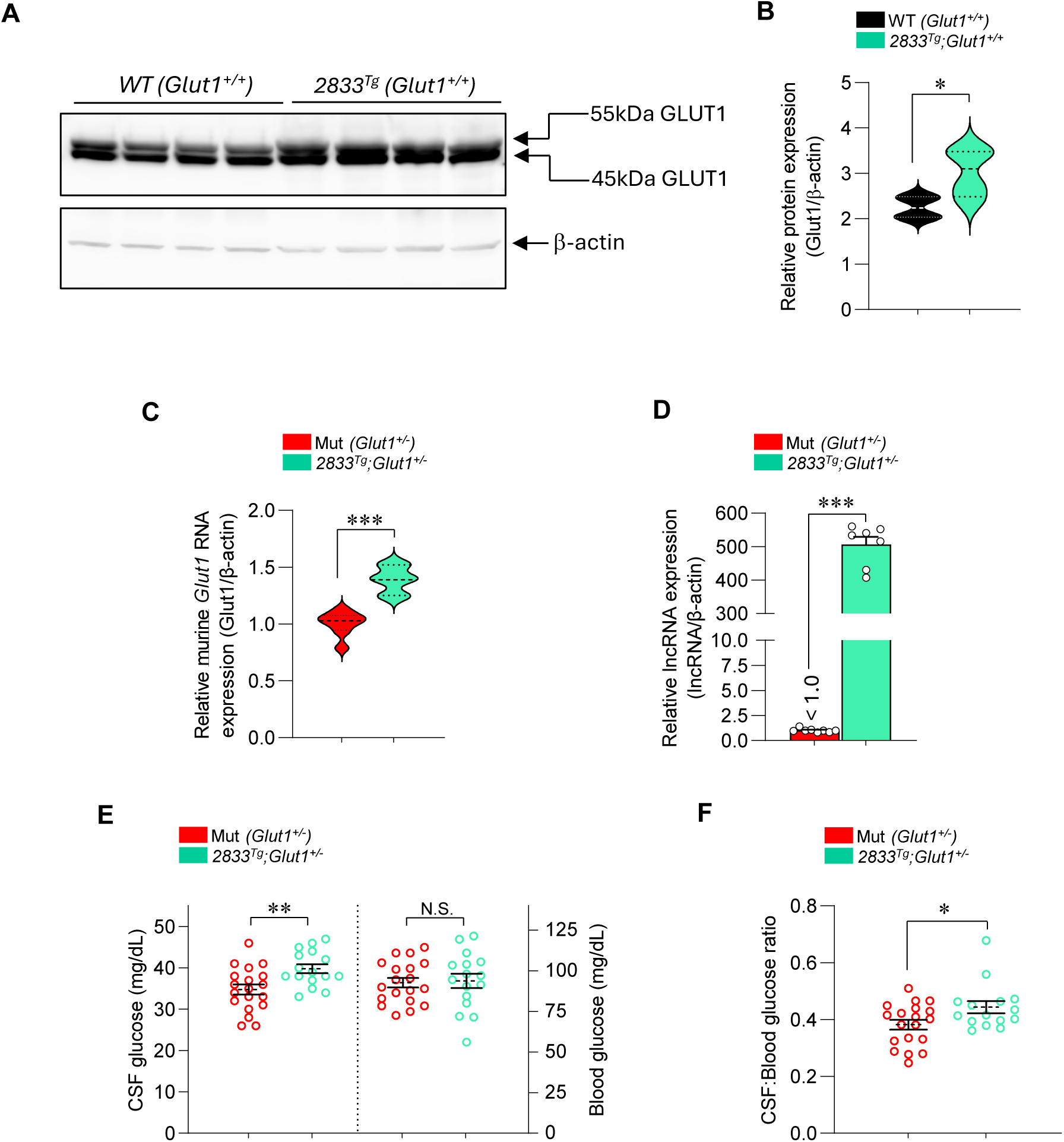
Transgenic expression of the human *SLC2A1-DT* lncRNA stimulates murine *Slc2a1 (Glut1)* expression. **(A)** Representative western blot depicting an increase in *murine* GLUT1 protein in brain tissue of transgenic mice harboring the intact human *SLC2A1-DT* lncRNA. Levels of protein in WT (non-transgenic) mice devoid of the lncRNA are shown for comparison. Note: 55kDa band – endothelial GLUT1; 45kDa band – astrocytic GLUT1. **(B)** Quantified levels of GLUT1 protein in the two sets of mice based on the blot in panel A. Note: *, *P* < 0.05, *t* test, n = 4 mice of each genotype. Graphs depicting quantified levels of **(C)** murine *Slc2a1 (Glut1)* transcripts and **(D)** *SLC2A1-DT* lncRNA in brain tissue of GLUT1DS mutants with or without the *SLC2A1-DT* lncRNA-containing transgene. Note: ***, *P* < 0.001, *t* test, n ≥ 7 subjects of each genotype for each of the assessments. **(E)** Graphical representation of an increase in CSF but not blood glucose in GLUT1DS mutants expressing the human *SLC2A1-DT* lncRNA transgene. **(F)** CSF:Blood glucose levels also increase in GLUT1DS mutants expressing the human *SLC2A1-DT* lncRNA transgene. Note: *, **, *P* < 0.05, *P* < 0.01 *t* tests, n ≥ 15 mutants of each genotype for the assessments in panels E, F.

**Supplemental Figure 2.**
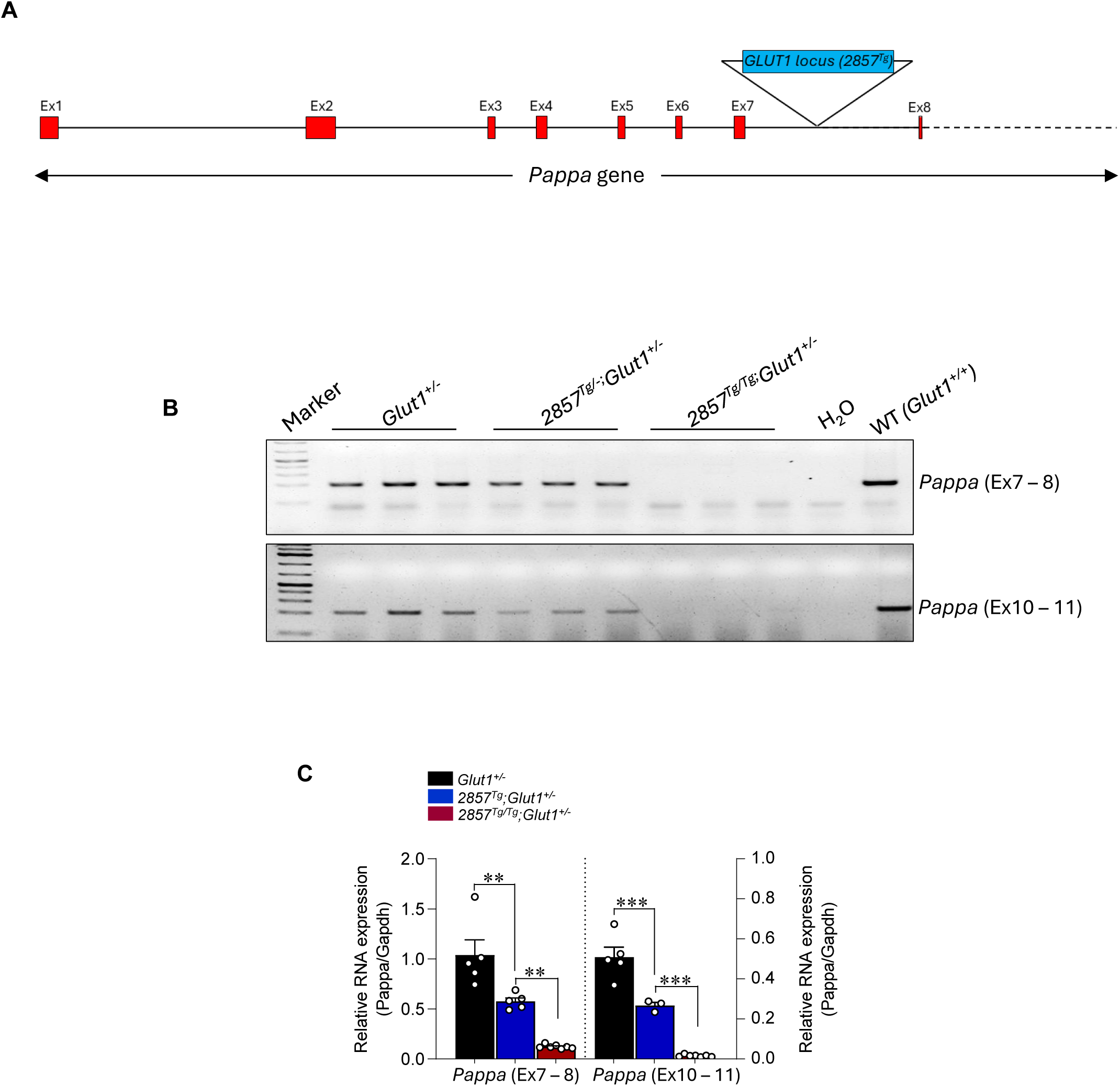
The 2857 *GLUT1* transgene is inserted into the *Pappa* gene rendering it a functional null. **(A)** Cartoon depicting the murine *Pappa* gene and the insertion site of the 2857 transgene in intron 7 of the gene. **(B)** Representative gel following RT-PCR of the *Pappa* gene in *Glut1^+/-^* mutants homozygous, heterozygous or devoid of the 2857 *GLUT1* transgene. Note absence of the *Pappa* transcript in mice homozygous for the 2857 transgene. **(C)** Quantified levels of the *Pappa* transcript, by Q-PCR, in the three cohorts of mice assessed in the previous panel. Note: **, ***, *P* < 0.01, *P* < 0.001, one-way ANOVA, n = 3 – 7 mutants of each genotype for the assessments.

